# Comparative Genomics of two Inbred Lines of the Potato Cyst Nematode *Globodera rostochiensis* reveals disparate Effector Family-specific Diversification Patterns

**DOI:** 10.1101/2021.03.15.435409

**Authors:** Joris J.M. van Steenbrugge, Sven van den Elsen, Martijn Holterman, Mark G. Sterken, Peter Thorpe, Aska Goverse, Geert Smant, Johannes Helder

## Abstract

**Background:** Potato cyst nematodes belong to the most harmful pathogens in potato, and durable management of these soil-borne parasites largely depends on host-plant resistances. These resistances are pathotype specific. The current *Globodera rostochiensis* pathotype scheme that defines five pathotypes (Ro1 - Ro5) is for fundamental and practical reasons barely useful. As a result, resistant potato varieties are worldwide used in a non-informed manner.

**Results:** We generated two novel reference genomes of *G. rostochiensis* inbred lines derived from a Ro1 and a Ro5 population. These genome sequences comprise 173 and 189 scaffolds respectively, marking a ≈ 24-fold reduction in fragmentation as compared to the current reference genome. We provide copy number variations for 18 effector families. Four dorsal gland effector families were investigated in more detail. SPRYSECs, known to be implicated in immune suppression, constitute by far the most diversified family with 60 and 99 variants in Ro1 and Ro5 distributed over 18 and 26 scaffolds. In contrast, CLEs, effectors involved in feeding site induction, show strong physical clustering. The 10 and 16 variants cluster on respectively 2 and 1 scaffolds. Given that pathotypes are defined by their effectoromes, we pinpoint the disparate nature of the contributing effector families in terms of sequence diversification and loss and gain of variants.

**Conclusion:** Two novel reference genomes allow for nearly complete inventories of effector diversification and physical organisation within and between pathotypes. Combined with insights we provide on effector family-specific diversification patterns, this constitutes a solid basis for an effectorome-based virulence scheme for this notorious pathogen.

## Introduction

Plant-parasitic nematodes have a significant impact on food and feed production worldwide. Every cultivated crop is parasitized by at least one nematode species, resulting in a net loss of over 70 billion US dollar annually (Nicol *et al.*, 2011). From an economic point of view root-knot and cyst nematodes have the highest impact (Jones *et al.*, 2013). Whereas root-knot nematodes have a higher impact in warmer climate zones, cyst nematode problems mostly occur in the temperate regions. Unlike root-knot nematodes, most cyst nematodes have a defined center of origin. For example, soybean cyst nematodes originate from north-east Asia and have spread as a successful and highly harmful parasite to all major soybean-growing areas. Potato cyst nematodes diversified in the Andes in South America, and have now proliferated to all major potato production areas in the world (*e.g*. Plantard *et al.*, 2008). Outside of their centers of origin, cyst nematodes belong to the most harmful pathogens of the crops mentioned above.

One of the most widely applied control measures is the use of resistant host plants. Resistances against potato cyst nematodes tend to have a long agronomic life span due to cyst nematodes’ unique biology. Potato cyst nematodes usually have only one generation per year through obligate sexual reproduction, go into diapause for months, and - once hatched - their motility is in the range of a few cm per day. These characteristics drastically slow down the process of selection and proliferation of virulent individuals. Potato breeders have introgressed the resistance gene *H1* from *Solanum tuberosum* ssp. *andigena* CPC 1674 into numerous potato cultivars. The *H1* gene confers resistance against *G. rostochiensis* pathotypes Ro1 and Ro4 (Toxopeus & Huijsman, 1952). A particularity of most of the dominant resistance (R) proteins is that they are effective against a subset of pathotypes within a species only.

Based on a number of differentials, pathotypes have been defined within the two potato cyst nematode species *G. rostochiensis* and *G. pallida.* Five pathotypes named Ro1 – Ro5 have been proposed for *G. rostochiensis*, whereas three pathotypes (Pa1, Pa2, Pa3) were discriminated within *G. pallida* (Kort *et al.*, 1977). Apart from being laborious and time-consuming, the current pathotype scheme has limited value as it lacks a solid genetic basis. The distinction between for instance the *G. pallida* pathotypes Pa2 and Pa3 is elusive (Phillips & Trudgill, 1983). For *G. rostochiensis*, genome-wide allele frequencies correlate with the geographical distribution of populations, regardless of pathotype (Mimee *et al.*, 2015; Thevenoux *et al.*, 2020). This indicates that the genetic basis of the pre-defined pathotypes is small. A robust pathotyping scheme for potato cyst nematodes is highly desirable because it would lead to far more efficient and durable use of the limited number of host plant resistances currently available in potato. The availability of high-quality reference genome sequences from individual pathotypes would be an ideal starting point for pathotypes’ molecular characterization.

Resistant plant species deploy R proteins as surveillance molecules that recognize specific effector molecules - or their activities - secreted by nematodes (refs). Nematodes use a protrusible stylet to inject effector proteins into plant cells. Effectors are diverse and fulfil functions ranging from plant cell wall degradation to the induction of a feeding site and suppressing the plant’s innate immune system (Ali *et al.*, 2015). The nematode produces effectors mainly in the subventral and the dorsal esophageal glands. Effectors are usually members of diversified gene families, and potato cyst nematode typically produces multiple variants per effector. An example is the SPRYSEC gene family that codes for a highly expanded set of proteins that act as activators and suppressors of plant defense (Diaz-Granados *et al.*, 2016). One variant of this family, RBP-1, was shown to trigger the activation of the potato resistance gene *Gpa2* (Sacco *et al.*, 2009) resulting in local hypersensitive response. Effector proteins secreted by the cyst nematode parasite are most likely responsible for the activation of plant resistance proteins. However, this was demonstrated for only a small number of resistance genes (*Gpa2;* Sacco et al., 2009*, Cf-2;* Lozano-Torr*es et al., 2012*).

Sequencing the genome of plant-parasitic nematodes is more challenging than for other, larger organisms. With the currently available methods, it is practically impossible to isolate and sequence DNA from an individual nematode to gain enough coverage to generate a high-quality reference genome sequence—especially when isolating high molecular weight DNA required for long-read sequencing technologies. Reference genomes of plant-parasitic nematodes are therefore often based on the genetic material from a population. Consequently, the reference genome includes a substantial heterozygosity level, as the starting material includes a high degree of allelic variation. The current reference genome sequences of potato cyst nematodes *Globodera rostochiensis* (Eves-van den Akker et al., 2016) and *G. pallida* (Cotton et al., 2014) were each generated using heterozygous starting material (selected field populations), and are relatively fragmented (respectively 4,377 and 6,873 scaffolds). In *G. rostochiensis*, Eves-van den Akker et al. (2016) predicted 138 high confidence effector genes based on sequence similarity with previously described effector gene families. Furthermore, a third of these genes were identified to cluster on effector gene islands. Among these expanded gene families, sequence divergence between different pathotypes was estimated as well. While many single nucleotide polymorphisms and insertions/deletions were observed (Eves-van den Akker *et al.*, 2016), the highly fragmented reference genome sequence made it challenging to distinguish between sequence and copy number variation. Less fragmentation in the genome sequence would similarly make it possible to display the degree of clustering of effector genes more accurately.

We generated a new set of reference genome sequences to allow for the accurate organization of effector genes and to compare copy number variation and sequence variation between the Ro1 and Ro5 pathotypes. A precise representation of these two sources of genetic variation is essential for developing molecular pathotyping methods in the future. The current reference genome sequence of *G. rostochiensis* shows a diploid genome size of 95.9Mb in size (Eves-van den Akker *et al.*, 2016) and is expected to spread over eighteen diploid chromosomes (Rouppe van der Voort *et al.*, 1996). For this, we used two *G. rostochiensis* lines, one fully avirulent and one fully virulent with regard to the *H1* gene. The starting materials for these lines were two distinct field populations sampled from The Netherlands (Ro1-Mierenbos) and Germany (Ro5-Hamerz) by Janssen *et al.* (1990). The selection process started with a single cross between an individual male and a female. After multiple generations, fully avirulent Ro1 (Gr-line19) and fully virulent Ro5 (Gr-line22) lines were generated regarding the *H1* resistance in potato (Janssen *et al.*, 1990). As a result, both Gr-Line19 and Gr-Line22 harbour limited genetic variation, with a theoretical maximum of 4 alleles per locus. For sexually reproducing species, this is the minimum level of heterozygosity that can be present in a population.

New genome assemblies were generated for each of the inbred lines based on PacBio long read-sequencing technology. Using these newly generated *G. rostochiensis* reference genome sequences with a substantially reduced number of scaffolds, we investigated the genomic organisation and the diversification of 18 effector families. A large number of differences in the number of paralogs and variation in sequence content were identified between the effector arsenals of the avirulent Gr-line19 and the virulent Gr-line22. These pathotype-specific effector variants form the basis for the generation of a virulence scheme for potato cyst nematodes.

## Methods

### DNA isolation and sequencing

Cysts from two *G. rostochiensis* lines that were selected for being fully avirulent Ro1 (Gr-line19) or fully virulent Ro5 (Gr-line22) with regard to the *H1* gene (Janssen *et al.*, 1990) were used as starting material for the collection of pre-parasitic second-stage juveniles (J2). J2’s were concentrated, and sucrose centrifugation was used to purify the nematode suspension (Jenkins, 1964). After multiple rounds of washing of the purified nematode suspension in 0.1 M NaCl, nematodes were resuspended in sterilized MQ water. Juveniles were lysed in a standard nematode lysis buffer with proteinase K and beta-mercaptoethanol at 60°C for 1 h as described in Holterman *et al.* (2006). The lysate was mixed with an equal volume of phenol: chloroform: isoamyl alcohol (25:24:1) (pH 8.0) following a standard DNA purification procedure, and finally, DNA was precipitated with isopropanol. After washing the DNA pellet with 70% ethanol for several times, it was resuspended in 10mM Tris-HCL (pH 8.0). DNAs of both inbred lines (each 10 −20 μg) were sequenced using Pacific Biosciences SMRT sequencing technology at Bioscience (Wageningen Research, Wageningen, The Netherlands) Gr-line19 was sequenced to a depth of approximately 119X with an average read length of 5,641 bp, whereas Gr-line22 was sequenced 132X with an average read length of 7,469 bp. Depth was calculated based on the assembly sizes. In parallel, a 2x 250 bp Illumina NovaSeq run resulted in 188x coverage of paired-end reads per line used to polish the initial assemblies. The raw sequencing reads and the genome assemblies are available under NCBI accession PRJNA695196.

### Genome assembly

Raw PacBio reads were first corrected by merging haplotypes with the correction mode of Canu v1.8 (Koren *et al.*, 2017), allowing a corrected error rate of 15% and a corrected coverage of 200. Using long-read assembler wtdgb2 v2.3 (Ruan & Li, 2020), approximately one hundred assemblies were generated per inbred line, optimizing the parameters minimal read length, k-mer size, and minimal read depth. The quality of the initial assemblies was assessed based on whether the assembly size was close to the genome size estimate (Eves-van den Akker *et al.*, 2016). Completeness of the genome was assessed using BUSCO v3 (Seppey *et al.*, 2019) using the standard library of eukaryotic single copy genes. Based on the criteria mentioned before, the most optimal assembly was then selected for each line and used for post-assembly processing.

After determining the most optimal assembly, remaining unmerged haplotigs were filtered from the assembly using Purge Haplotigs v1.0.4 (Roach *et al.*, 2018). The assembly was then tested for contamination using the blobtools pipeline v.1.0.1. Contigs were scaffolded with PacBio reads using SSPACE-Longread (Boetzer & Pirovano, 2014). The remaining gaps in the scaffolds were then filled using a consensus alignment approach. NovaSeq data were used to polish the resulting assemblies using three iterations of Arrow v2.3.3 and five iterations of Pilon v1.23 each. Repeat regions were soft masked using RepeatModeler v1.0.11 and RepeatMasker v4.0.9. Gene annotations were predicted for both assemblies using BRAKER v2.1.2 aided by RNAseq datasets of different *G. rostochiensis* life stages that were mapped using Hisat v2.2.0 (NCBI BioProject accessions: PRJEB12075, PRJNA274143, PRJNA274143).

### Genome Synteny

Genome synteny was determined between the genome assembly of Line19, Line22 and the previous Ro1 reference genome (NCBI BioProject PRJEB13504) through a progressive genome alignment using Mauve v2.4.0. The alignment was then visualized in Circos v0.69-9, showing only syntenic regions of 3kb and larger.

### Estimating Heterozygosity levels and structural variation

Heterozygosity levels were estimated based on the frequency of heterozygous versus homozygous variants (SNPs and small indels). A comparison was made between Gr-Line19 and the JHI-Ro1 population, using the Gr-Line22 genome assembly as a reference. Illumina reads were mapped with Burrows-Wheeler Aligner (BWA) using default settings. For Gr-Line19, a library of Illumina NovaSeq reads (Accessions: SRR13560389, SRR13560388) was used, and for JHI-Ro1, Illumina HiSeq reads (Accessions: ERR114519) were mapped against the reference. Variants were called with bcftools v.1.9 with multiallelic variant calling enabled, at a maximum depth of 1,000 reads.

The structural variation between the newly generated assemblies of Gr-Line19 and Gr-Line22 was estimated by the frequency of heterozygous *versus* homozygous structural variants (SVs) with fragment size > 1kb. Raw Pacbio reads of Gr-Line19 were mapped against the Gr-Line22 (and *vice versa*) using NGMLR v0.2.7 with default settings. SVs were then called using Sniffles v1.0.10 running the standard settings (Sedlazeck *et al.*, 2018).

### Identification of Effector Homologs

Effector genes were identified in both line Gr-line19 and Gr-line22 based on the proteomes predicted by BRAKER2. Phobius (Käll *et al.*, 2004) was used to check for the presence of a signal peptide for secretion. Homologs for glycoside hydrolase (GH) families 5, 30, 43, 53, Pectate lyase 3, Glutathione Synthetase were identified with HMMER v3.2.1 based on pre-calculated profile HMMs in the PFAM database (entries PF00150, PF02055, PF04616 and PF07745, PF03211, PF02955 respectively). SPRYSEC homologs were identified by testing protein sequences for a SPRY domain (hmm profile PF00622). Arabinogalactan galactosidase homologs were identified with a custom profile HMM-based on UniProt sequences (entries O07012, Q65CX5, Q65CX4, D9SM34, P48841, O31529, Q8X168, Q5B153, O07013, P83692, P48842, P83691, Q4WJ80, B0XPR3, A1D3T4, Q2UN61, Q0CTQ7, A2RB93, Q9Y7F8, B8NNI2, Q76FP5). CLE-like homologs were identified with a custom profile HMM-based on UniProt sequences (D1FNJ7, D1FNK5, D1FNJ9, D1FNK2, D1FNK8, D1FNK3, D1FNK0, D1FNK4). GenBank peptide sequences JQ912480 to JQ912513 were used to generate a custom profile HMM for the effector family 1106. Based on GenBank entries KM206198 to KM206272, a custom profile HMM was made for the HYP effector family. Homologs of the *Hetereodera glycines* effector families Hg16B09 (GenBank: AAO85454) and GLAND1-18 (GenBank: KJ825712 to KJ825729) were identified with blastp, with the following cut-offs: an identity score higher than 35%, a query coverage of at least 50%, and an E-value lower than 0.0001

### Phylogeny

Multiple Sequence Alignments were generated based on the coding sequences of the orthologs per effector family, using Muscle v3.8.1551 using standard options. Phylogenetic trees were then generated with RaxML v8.2.12 running a GTRGAMMA model with 100 bootstrap replicates. The resulting trees were visualized and organized in Figtree (v. 1.4.4).

## Results

### Genome Assemblies

Two inbred lines of the potato cyst nematode *G. rostochiensis* were initially derived from crossings between individuals from two populations, Ro1-Mierenbos and Ro5-Harmerz (Janssen *et al.*, 1990). DNA from these lines, Gr-Line19 and Gr-line22, were sequenced using PacBio sequencing technology with respectively 119X and 132X coverage and assembled into two reference genome sequences (Table 1). Benefitting from this long read technology and the significantly smaller genetic background, the two newly generated *G. rostochiensis* genome assemblies are less fragmented than the first genome sequence that was published (nGr.v1.0) (Eves-van den Akker *et al.*, 2016) while maintaining a comparable assembly size. The number of scaffolds in the new assemblies is about 24 fold lower than in the original *G. rostochiensis* reference genome sequence (Table 1). At the same time, the scaffold N50 increased about 20 fold from 0.085 to around 1.7 Mb. Regarding the assembly size and BUSCO score, the novel assemblies are comparable to the current reference. The assemblies of Gr-Line19 and Gr-Line22 harbor 2,733 and 6,572 gaps, respectively, covering in total 130Kb and 150Kb. As compared to the current *G. rostochiensis* reference (Eves-van den Akker *et al.*, 2016), the number and lengths of gaps showed a 29-fold reduction.

**Table 1.** Comparative genome statistics of three *G. rostochiensis* genome assemblies

| *G. rostochiensis* population | Assembly ID | Size (Mb) | # scaffolds | Scaffold N50 (Mb) | BUSCO notation assessment results (in %) |  |  |  |
| --- | --- | --- | --- | --- | --- | --- | --- | --- |
|  |  |  |  |  | Single | duplicated | fragmented | missing |
| Ro1 population from JHI PCN collection (JHI-Ro1) | nGr.v1.0 | 96 | 4,377 | 0.085 | 82.8 | 1.0 | 8.3 | 7.9 |
| Gr-Line19 | Gr19v10 | 92 | 173 | 1.70 | 82.2 | 1.7 | 7.9 | 8.2 |
| Gr-Line22 | Gr22v10 | 101 | 189 | 1.80 | 81.5 | 1.3 | 8.3 | 8.9 |

BUSCO (Benchmarking Universal Single-Copy Orthologs) - eukaryota\_odb10. For JHI-Ro1 nGr.v1.0 see Eves-van den Akker *et al.* 2016

The repeat content in both reference genome sequences is relatively low, 2.6% for Gr-Line19 and 1.6% for Gr-Line22. The GC content in repeat regions for Gr-Line19 (40.3%) was comparable to this genotype’s overall GC content (39.1%). In Gr-Line22, the GC content in repeat regions (32.5%) was lower than the overall GC content (38.3%). In predicted protein-coding regions, the GC content is comparable between both reference genome sequences (Gr-Line19: 50.8%, Gr-Line22: 50.9%). Using Braker2 as a gene-prediction tool, 17,928 and 18,258 genes were predicted in the Gr-Line19 and Gr-Line22 genome assemblies, coded for 21,037 21,514 transcripts, respectively. The protein-coding regions tak9e up approximately 33% (Line19) and 30% (Line22) of the genomes at an average density of 89.3 (Gr-Line19) and 86.6 (Gr-Line22) genes per Mb.

Synteny between the newly generated genomes and the current reference genome (Eves-van den Akker *et al.*, 2016) was evaluated using a progressive genome alignment. Homologous regions larger than 3 kb and their genomic organization are presented in Figure 1. A broad span of regions in the nGr.v1.0 reference assembly shows homology to both new assemblies (respectively 67%, 72%, and 61% of the total assembly sizes for JHI-Ro1, Gr-Line19 and Gr-Line22). While the total numbers of base pairs that are covered in a homologous region are roughly within a 10% range of each other, both new assemblies show substantially larger continuous and so far uncovered regions.

**Fig. 1.**
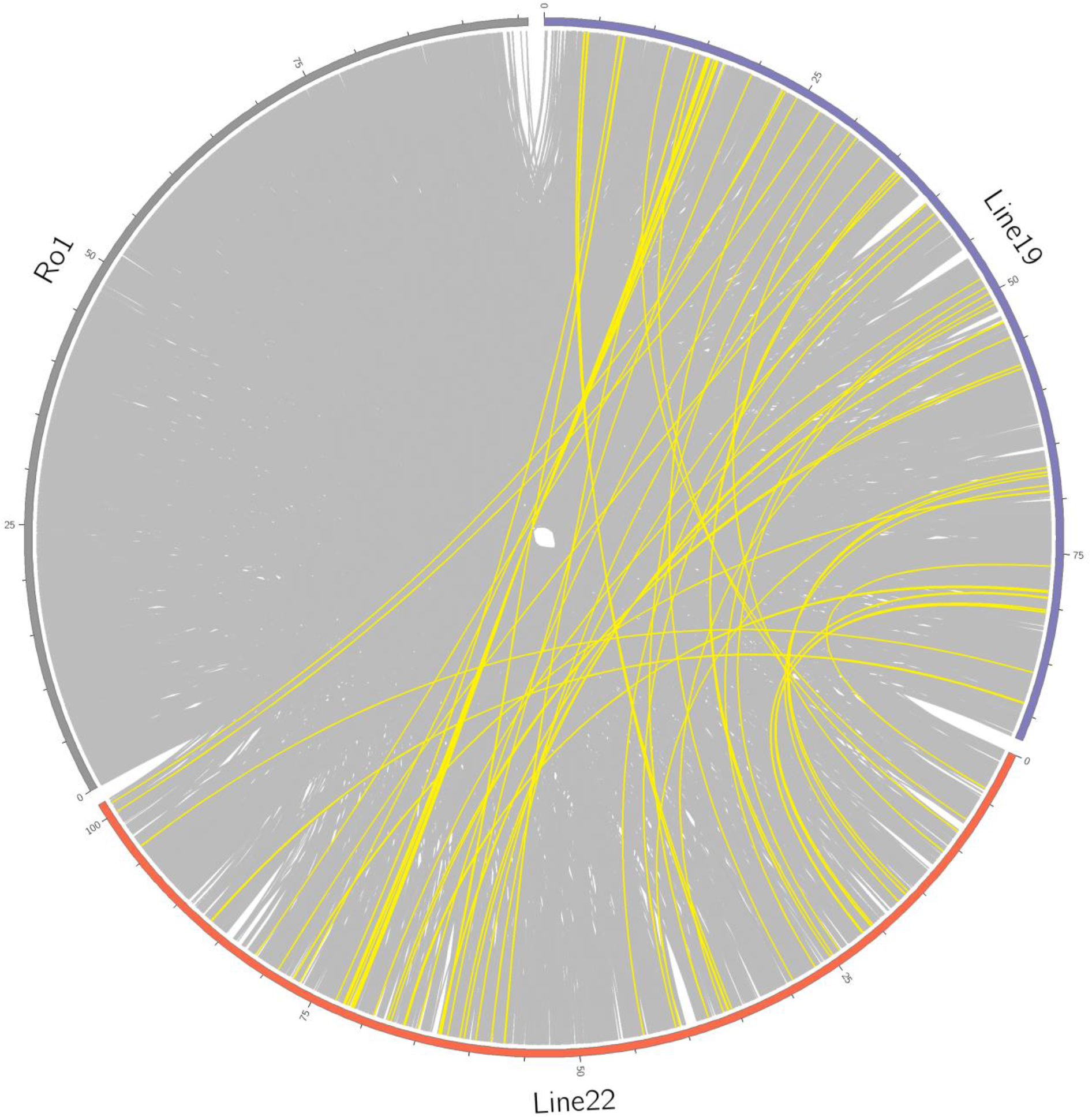
Synteny between Gr-Line19, Gr-Line 22, and JHI-Ro1. Only syntenic regions larger than 3 kb are shown. Yellow lines represent regions that are only present in Gr-Line19 and Gr-Line22.

### Heterozygosity and structural variation between the two *Globodera rostochiensis* genomes

The inbred lines Gr-Line19 and Gr-Line22 originate from single crossings of individuals, and, as a result, the genetic variation is expected to be smaller than for field populations. To pinpoint the effect of this genetic bottleneck caused by a single crossing, a comparison was made between the proportion of heterozygous and homozygous single nucleotide variants between Gr-Line19 (Ro1) and JHI-Ro1, the selected field population used to generate the current *G. rostochiensis* reference genome (Eves-van den Akker *et al.*, 2016) while using the Gr-Line22 (Ro5) genome as a reference. Among the called variants that passed the quality filter (JHI-Ro1 n=584,145; Gr-Line19 n=716,491), 37% of the JHI-Ro1 loci were homozygous, as compared to 47% of the variants in Gr-Line19 (Figure 2A). The increased level of homozygosity in Gr-Line19 reflects the relatively narrow genetic basis of this inbred line.

**Fig. 2.**
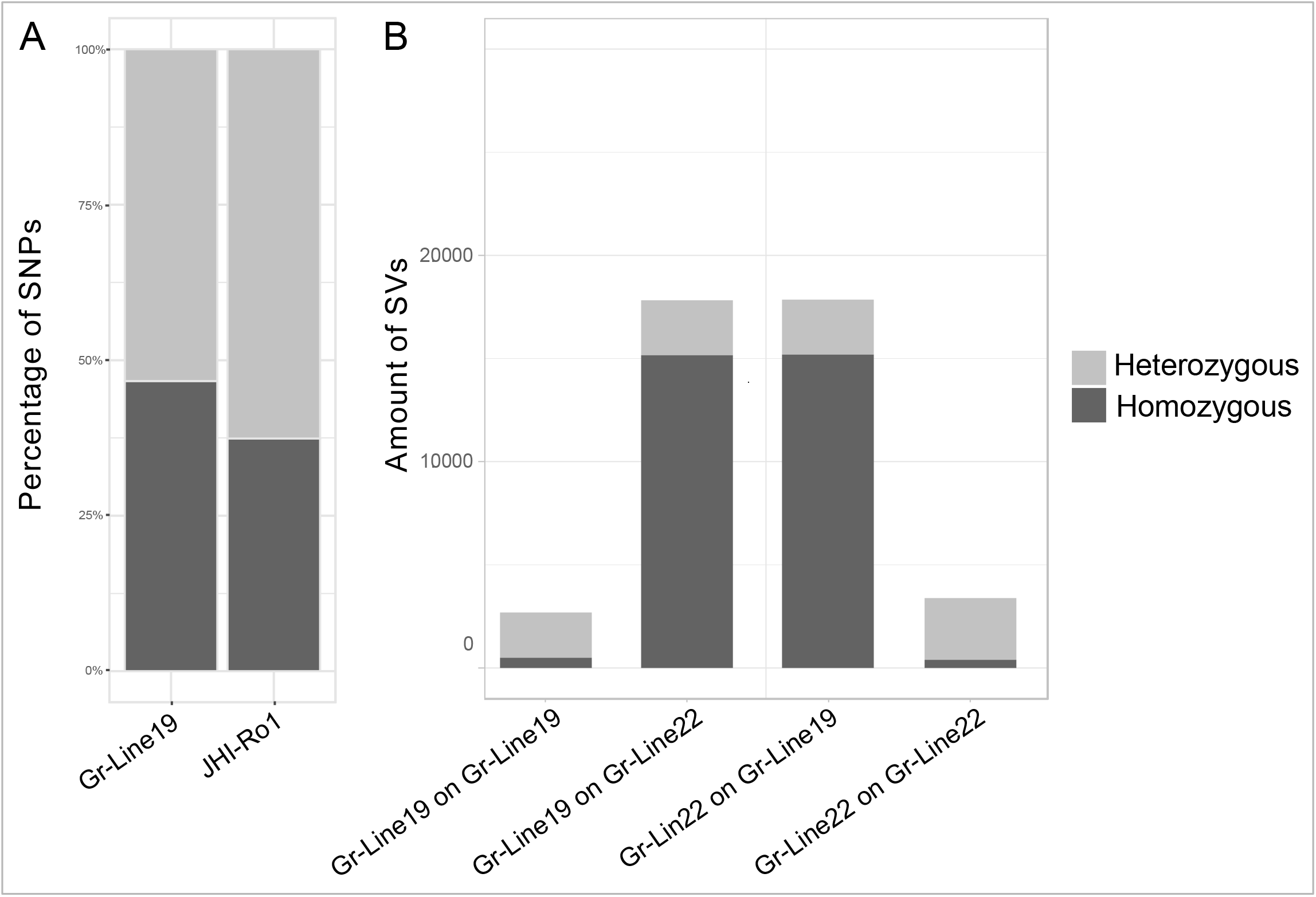
A. Relatively level of SNP heterozygosity of Gr-Line19 and JHI-Ro1 (JHI – James Hutton Institute, Scotland, UK). Both lines were compared to Gr-Line22. JHI Ro1 was used as the current *G. rostochiensis* reference genome. B. Comparison overall genomic constitution of *G. rostochiensis* inbred lines 19 and 22. In this overview structural variants are shown. Structural variants are DNA region of approximately 1 kb and larger in size, and can include inversions, insertions, deletions etc.

Secondly, structural variation (e.g. insertions, deletions, inversions) of approximately 1kb or larger within the individual lines and between Gr-Line19 and Gr-Line22 was determined. The proportions of heterozygous and homozygous structural variants with fragment sizes > 1kb were compared (Figure 2B). The structural variation within Gr-Line19 and Gr-Line22 was minimal (Figure 2B). This observation confirms the low level of structural intra-population heterozygosity. The proportion of homozygous variants was nearly identical while comparing Gr-Line19 with Gr-Line22 and *vice versa* (Gr-Line22 versus Gr-Line19: 85.09% & Gr-Line19 on Gr-Line22 85.06%).

### Expansion of Effector gene Families

We identified homologs of 18 known effector gene families from which at least one member was shown to be expressed in the subventral (6) or the dorsal (11) oesophageal gland cells, or in the amphids (1) (Table 2, Figure 3A).

**Table 2.**
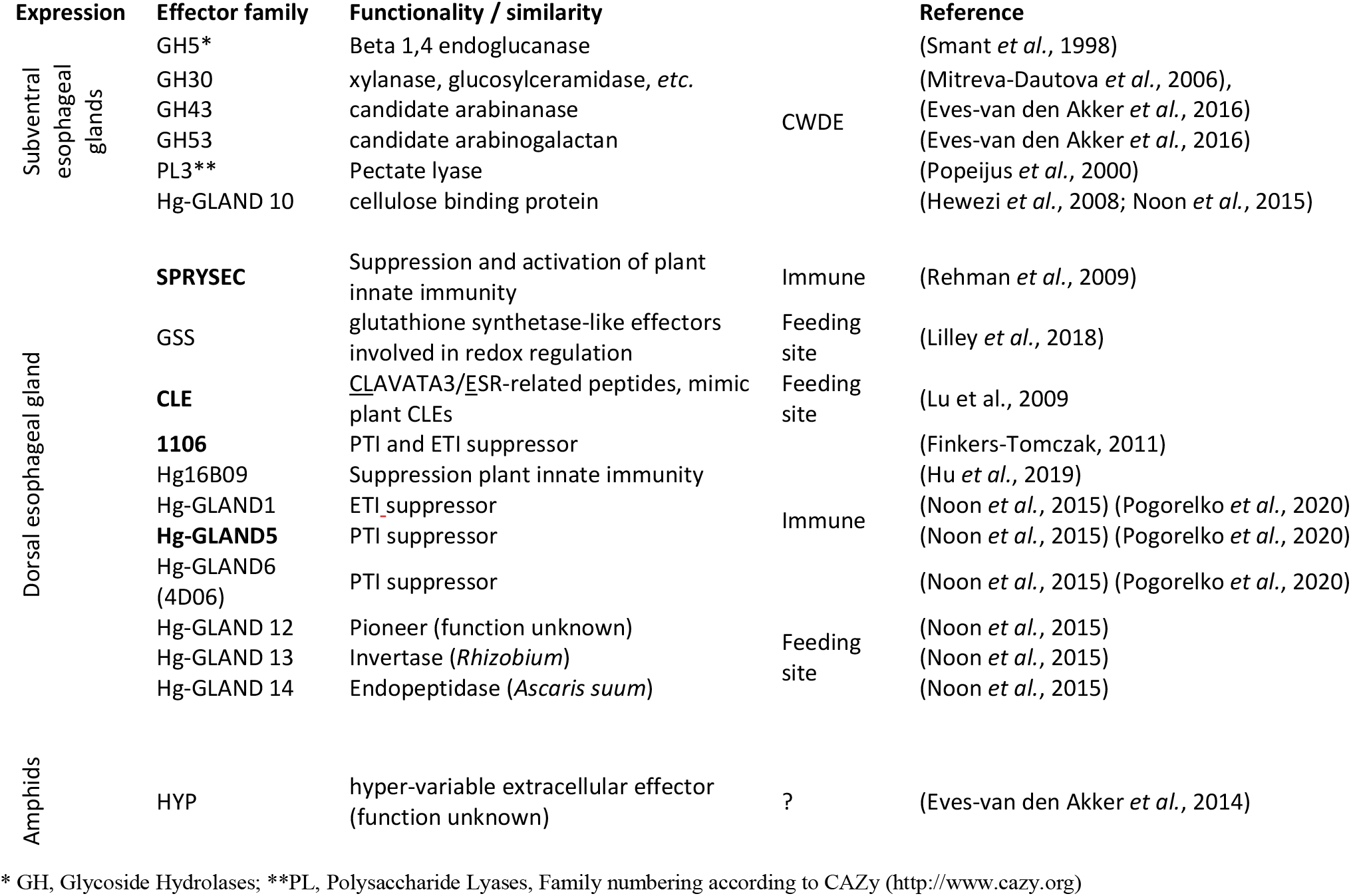
Effector families mapped in genomes of *G. rostochiensis* lines Gr-line19 and Gr-line22. Expression refers to nematode organs in which at least one effector family member was shown to be expressed. Subventral esophageal glands of potato cyst nematode are mainly active during migration to the host plant, and host plant penetration. The dorsal esophageal gland shows highest activity during feeding site induction and maintenance. Amphids are chemosensory organs located at the head region of the nematode. For effector families in bold diversification and physical distribution are investigated in detail. CWDE: cell wall-degrading enzymes, Immune: effector families for which at least one member is known to affect the plant innate immune system, Feeding site: effector families for which at least one member is known to be involved in feeding site induction or maintenance.

**Fig. 3.**
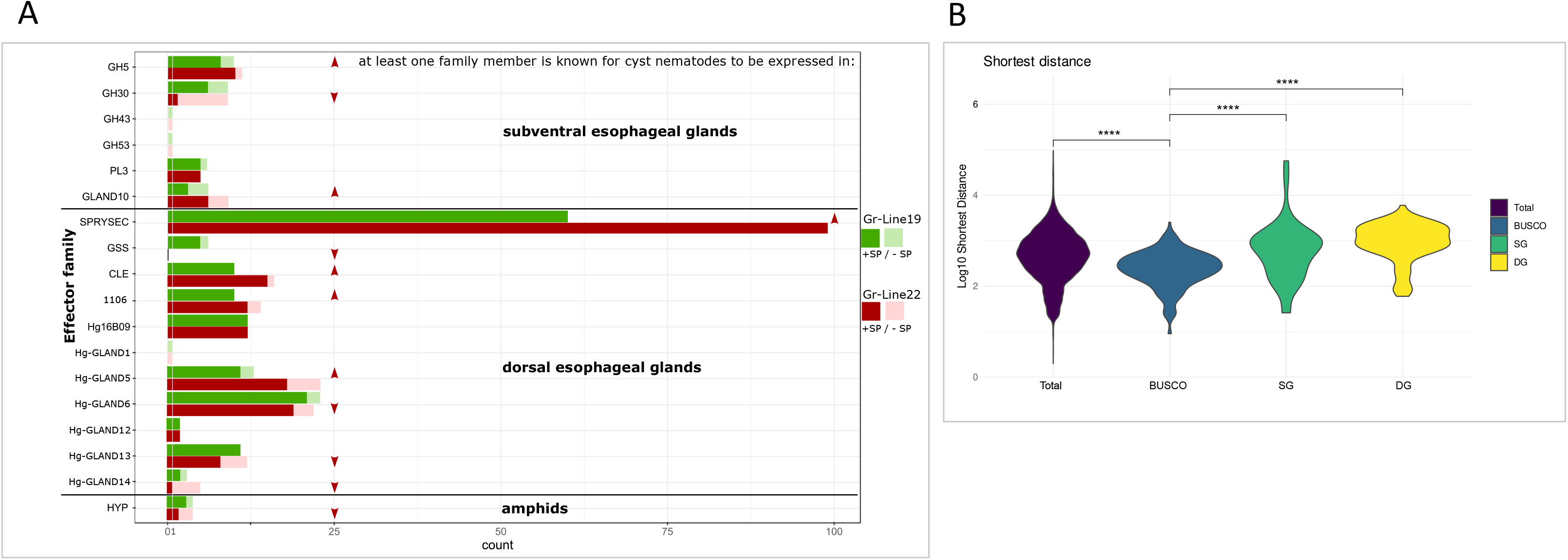
A. Copy numbers of effector gene families expressed in the secretory gland cells. A comparison was made between Gr-Line19 (Green) and Gr-Line22 (Red). Bright shades of green/red indicate copies that contain a signal peptide for secretion. Faded shades of green/red indicate the absence of a signal peptide for secretion. B Gene density comparison of all genes (total), eukaryotic universal single copy genes (BUSCO), effector genes secreted from the subventral esophageal gland cells (SG), and effector genes secreted from the dorsal esophageal gland cells (DG). The shortest distance of each gene was based on the closest adjacent gene (3’ or 5’) and is measured as the log10 number of basepairs. Statistical significance for each group was determined by comparison with the BUSCO gene set using a Wilcoxon test.

For each of these gene families, the copy number differences between Gr-Line19 and Gr-Line22 were determined. The number of paralogs per effector families varied from 99 SPRYSEC variants in Gr-Line22 to a single Hg-GLAND14 gene with signal peptide in the same line. Among the 18 effector families, six have a lower number of paralogs in Gr-Line22, six have an equal number of paralogs, whereas six have a higher number of variants in Gr-Line22 (Figure 3A).

Four effector families show a relatively large difference in the number of paralogs between Gr-Line19 and Gr-Line22. SPRYSEC is by far the most speciose effector family in both lines, but Gr-Line22 harbor 36 more variants with signal peptide than pathotype Ro1. Similarly, 11 Hg-GLAND5 homologs were present in Gr-Line19, while Gr-Line22 comprised 18 paralogs with a signal peptide. The reverse was also observed for Glutathione synthetase (GSS). Five copies were found in Gr-Line19, whereas this effector family is not present in Gr-Line22. Though less radical, a drastic decrease was observed for the subventral gland effector family GH30. Whereas six variants with signal peptide were identified in Gr-Line19, only two were found in Gr-Line22. It is noted that the GH30 family harbors various glycoside hydrolases that were previously categorized as GH5.

### Genomic organisation of effector genes

To characterise the genomic organisation of effector genes, the shortest distance between each gene and the closest adjacent gene was calculated (either at the 3’ or 5’-end of the full genomic sequence). This was done for effector genes, as well as for known non-effector genes (i.e. BUSCO gene set). The distances based on the full set of predicted genes ranged from extremely gene sparse to extremely gene dense regions (Figure 3B). BUSCO genes are generally located in regions that are more gene dense than expected at random (Wilcoxon Rank Sum test P < 0.0001). Effector genes expressed in either the dorsal or the subventral esophageal gland cells are often located in more gene sparse regions both as compared to non-effector genes and to any random gene (Wilcoxon Rank Sum test P < 0.0001).

Furthermore, the spatial organization and diversification between two pathotypes of *G. rostochiensis* lines is presented for four selected effector families that are expressed in the dorsal esophageal glands during parasitic life stages. Hg-GLAND5 effectors are known as plant triggered immunity suppressors (Pogorelko *et al.*, 2020). Members of the effector family 1106 were demonstrated to suppress both plant triggered immunity and effector triggered immunity (Finkers-Tomczak, 2011). The highly speciose SPRYSEC family was shown to be involved in both the suppression and the activation of the plant immune system (Ali *et al.*, 2015). CLE-like effectors were demonstrated to be involved in feeding site induction by mimicking the functionality of endogenous host-plant CLE peptides (Mitchum *et al.*, 2012).. Concentrating on the distribution of individual family members over the relevant scaffolds, large differences in the level of clustering per family are observed (Fig. 4). While the 60 and 99 SPRYSEC variants are distributed over respectively 18 and 26 scaffolds, the moderately diversified CLE family is concentrated on two scaffolds in case of Gr-Line19, and on a single scaffold for Gr-Line22.

**Fig. 4.**
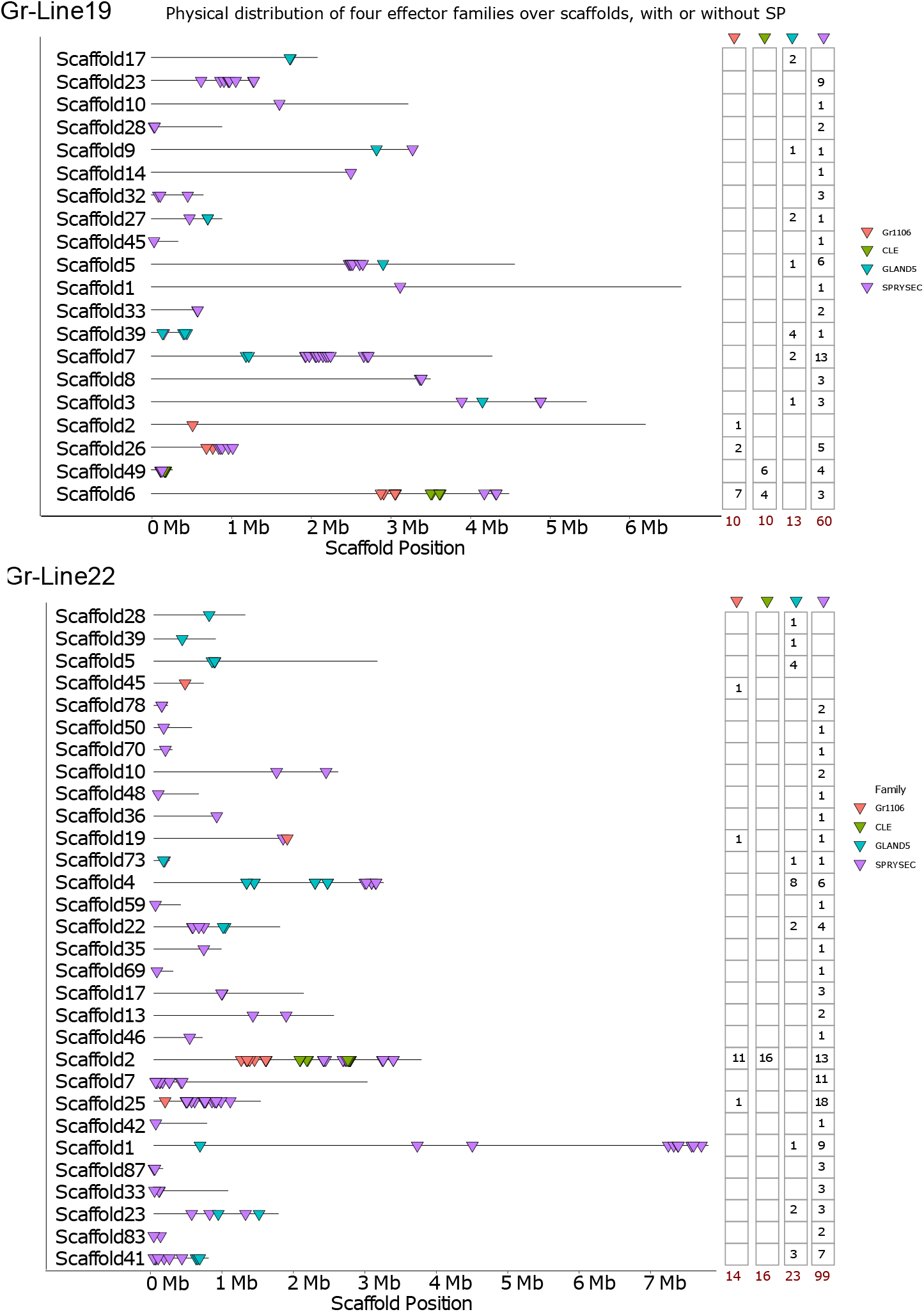
Spatial distribution of genes belonging to the effector families Gr1106 (Red), CLE (Green), GLAND 5 (blue), and SPRYSEC (purple). Each triangle indicates the genomic position of a single gene. At the right, the number of variants per effector family are given for each scaffold.

### A. Hg-GLAND5

Hg-GLAND5, also referred to as ‘putative gland protein G11A06’ (Gao et al., 2003), has first been discovered in soybean cyst nematode *Heterodera glycines*. This effector is expressed in the dorsal gland during a range of parasitic life stages, and it functions as a PTI suppressor (Pogorelko et al., 2020). In a transcriptional analysis of two *H. glycines* races, the expression level of Hg4J4-CT26, a GLAND5 family member, was shown to be highly race-dependent (Wang et al., 2014). Searches in public genome database revealed that both PCN species harbour homologs of HG-GLAND5 (Yang et al., 2019). In *G. rostochiensis*, the GLAND5 effector family comprises 13 and 23 members in Gr-Line19 and Gr-Line22, respectively. Among these variants, three and five are unlikely to be involved in parasitism as the corresponding protein sequences are not preceded by a signal peptide (SP) for secretion (Fig. 3A). A phylogenetic analysis of the GLAND5 family based on the coding sequences using RaxML revealed an initial split between four effectors without an SP on the one hand, and all functional GLAND5 effectors on the other (Fig. 5). Four clusters with mainly secreted GLAND5 variants could be discerned. Local expansion of effector variants in Gr-Line22 is present in three clusters. Most notable is the diversification in Box A, where eight related GLAND5 representatives from Gr-Line22 surround a single Gr-Line19 variant. Box C and Box D illustrate expansion in Gr-Line22 as well, although less extreme. Alternatively, Box B shows expansion in Gr-Line19, where two representatives of Gr-Line19 are present and a single variant of Gr-Line22.

**Fig. 5.**
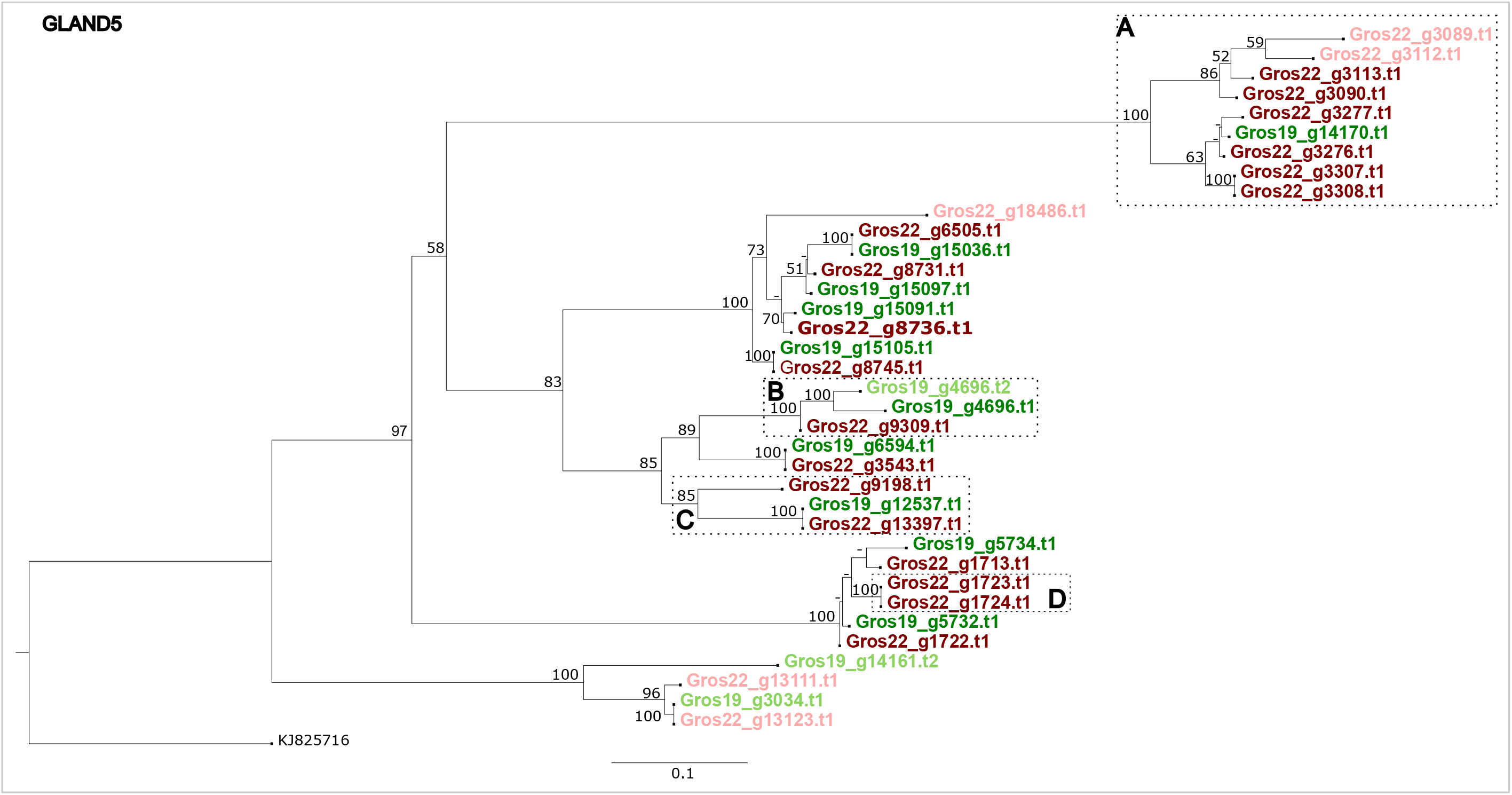
Phylogeny of GLAND5 effector genes of both Gr-Line19 (Green) and Gr-Line22 (dark red). A multiple sequence alignment was made using MUSCLE on the coding sequence. A phylogenetic tree was made using RAxML using a GTRGAMMA model, validated by 100 bootstrap replicates. Lighter shades of green or red indicate effector variants that lack a signal peptide for secretion.

### B. 1106

The 1106 gene family encodes mainly secreted proteins, and members were demonstrated to suppress bot PTI and ETI responses (Finkers-Tomczak, 2011). In this study, a conserved region of 1106 variants was shown to hybridize in the dorsal gland of infective juveniles of *G. rostochiensis*. Gr-Line19 contains ten paralogs, whereas 14 1106 paralogs were found in Gr-Line22. In terms of organization, the genes in Gr-Line22 and Gr-Line19 show a comparable degree of physical clustering (Fig. 4). To investigate the diversification of the effector family 1106, the phylogenetic relationship between the variants identified in Gr-Line 19 and Line 22 was examined (Fig. 6). In many cases, a 1106 variant in Gr-line 19 had a single, orthologous equivalent in Gr-line22 (see *e.g.* Gros19_2102 and Gros22_4744, and Gros19_2104 and Gros22_4746). Notably, the relationship between the small clusters of 1106 variants was largely unresolved. Within cluster A in Fig. 6, four Gr-Line22 variants were present, and only one representative from Gr-Line19. Two variants deviate substantially from most other 1106 family members. It is noted that these variants are not preceded by a signal peptide for secretion and thus are unlikely to act as effectors. Clusters B and C are highlighted as they might represent examples of subclades with an extra Line22-specific effector variant (Cluster B), or an additional Gr-Line19 variant (Cluster C). It should be noted, however, that the support for Cluster C is poor.

**Fig. 6.**
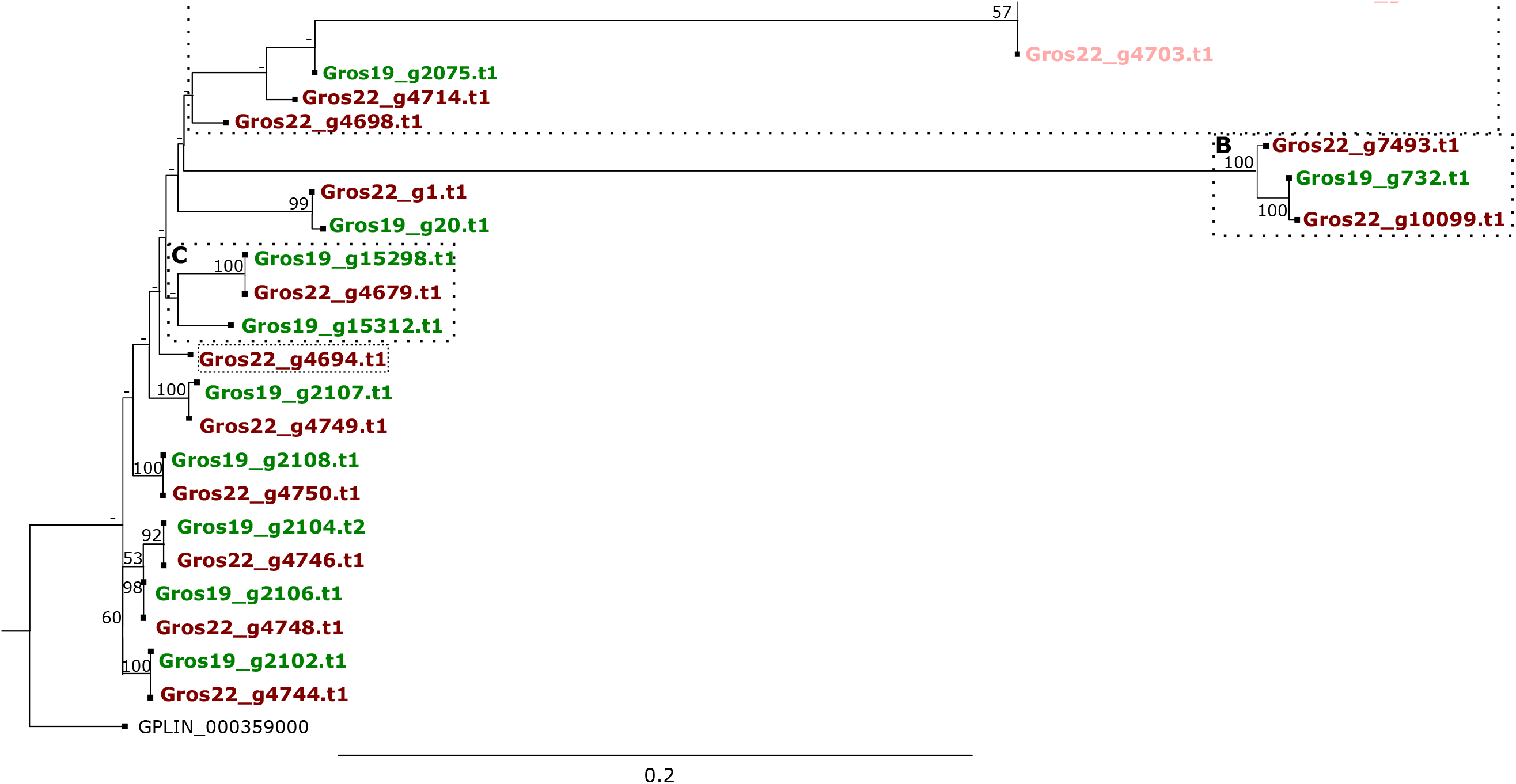
Phylogeny of Gr1106 effector genes of both Gr-Line19 (green) and Gr-Line22 (dark red). A multiple sequence alignment was made using MUSCLE on the coding sequence. A phylogenetic tree was made using RAxML using a GTRGAMMA model, validated by 100 bootstrap replicates. Lighter shades of green or red indicate effector variants that lack a signal peptide for secretion

### C. SPRYSEC

The SPRYSEC gene family encodes for secreted proteins that contain an <u>SP</u>la and <u>RY</u>anodine receptor. SPRYSECs are produced in the dorsal esophageal glands, and this effector family is thus far the most expanded one among the plant-parasitic nematodes (Diaz-Granados *et al.*, 2016). Several SPRYSECs from *G. rostochiensis* were shown to be implicated in the suppression and the activation of defence-related cell death (Postma *et al.*, 2012). Suppression was demonstrated for the variants SPRYSEC-4, −5, −8, −15, −18, and −19, whereas SPRYSEC15 elicited a defence response in tobacco (Ali *et al.*, 2015). In the closely related cyst nematode species *G. pallida,* a single SPRYSEC variant - RBP-1 - was shown to be responsible for the evasion of the potato resistance gene Gpa2, thus preventing a local HR (Sacco et al. 2009). No direct ortholog of RBP-1 could be found among the *G. rostochiensis* SPRYSECs (identity lower than 50%), with the used filtering criteria.

The diversification of the SPRYSEC-like variants in Gr-Line19 and Gr-Line22 was investigated by analysing the phylogenetic relationships. Although the number of SPRYSECs in Gr-Line19 (n=60) was already higher than for any other effector family, Gr-Line22 was shown to harbour even more members of this effector family (n = 99) (Fig. 3A). Maximum-likelihood-based inference revealed several SPRYSEC clusters (Fig. 7). Due to the poor backbone resolution, no statements can be made about the relationship between these clusters. It is noted that the support values for the more distal parts of the SPRYSEC tree are substantially higher than the support values for most of the more proximal bifurcations. Three large (A, B, and D) and two smaller (C, E) SPRYSEC clusters could be identified. The majority of Gr-Line19 variants have a single orthologous equivalent in Gr-Line22, while Gr-Line22 shows expansion in each of the clusters.

**Fig. 7.**
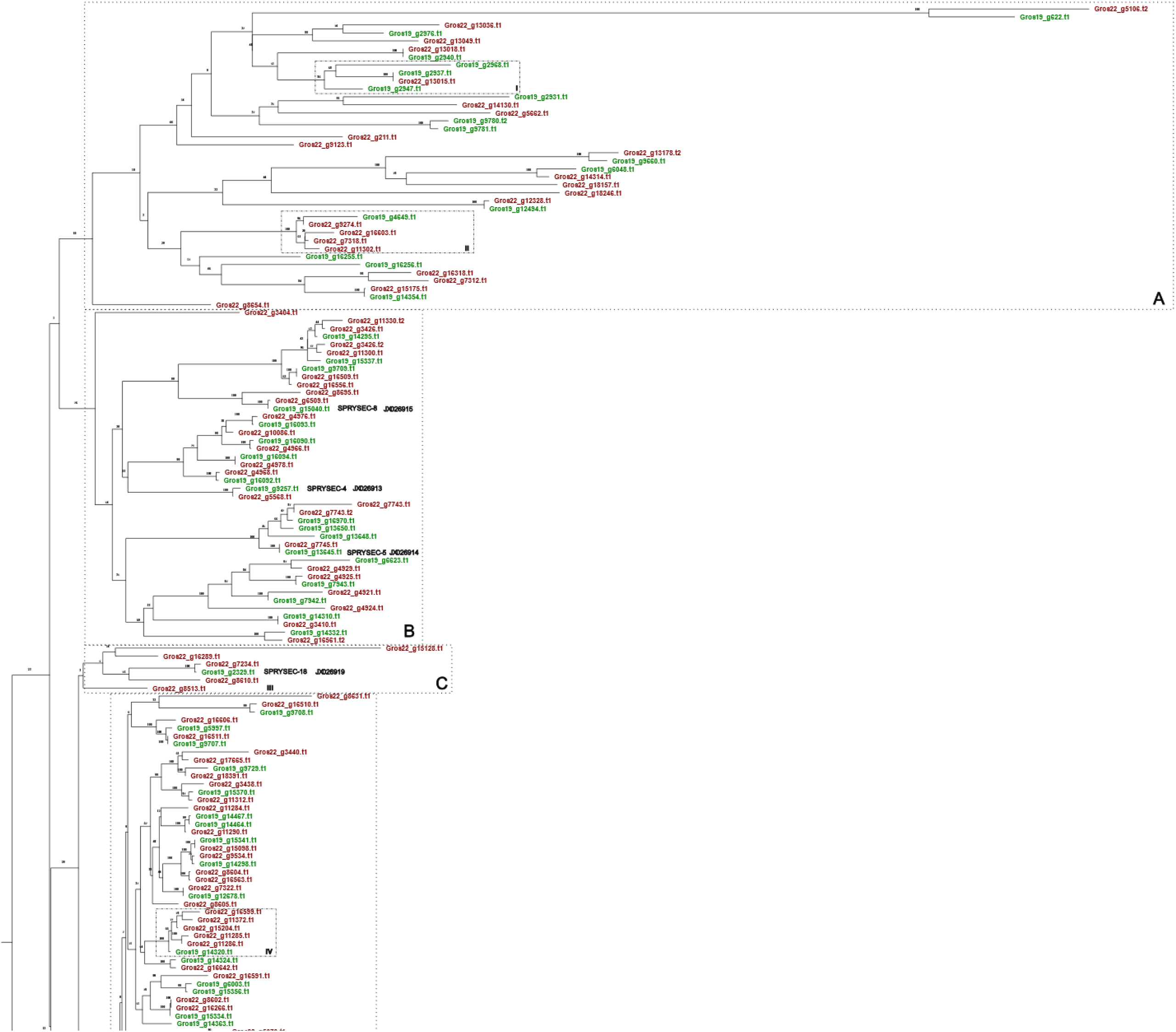
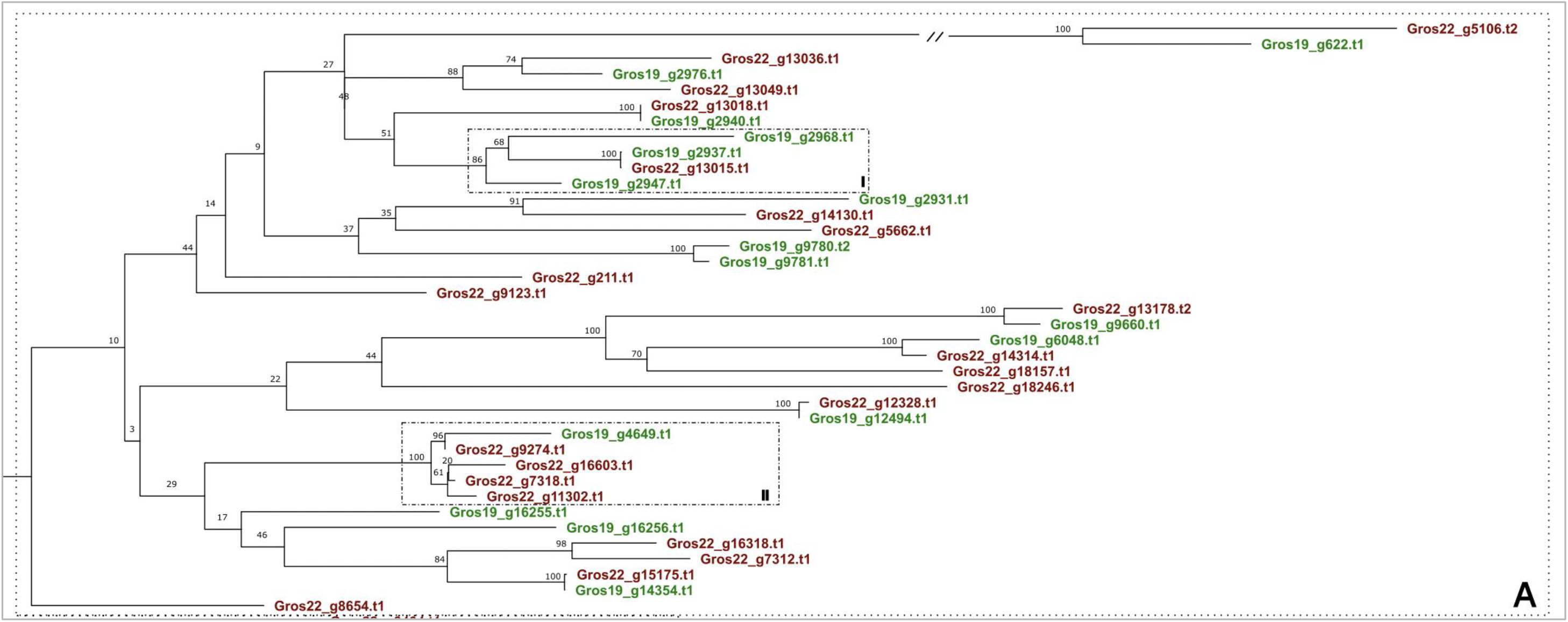
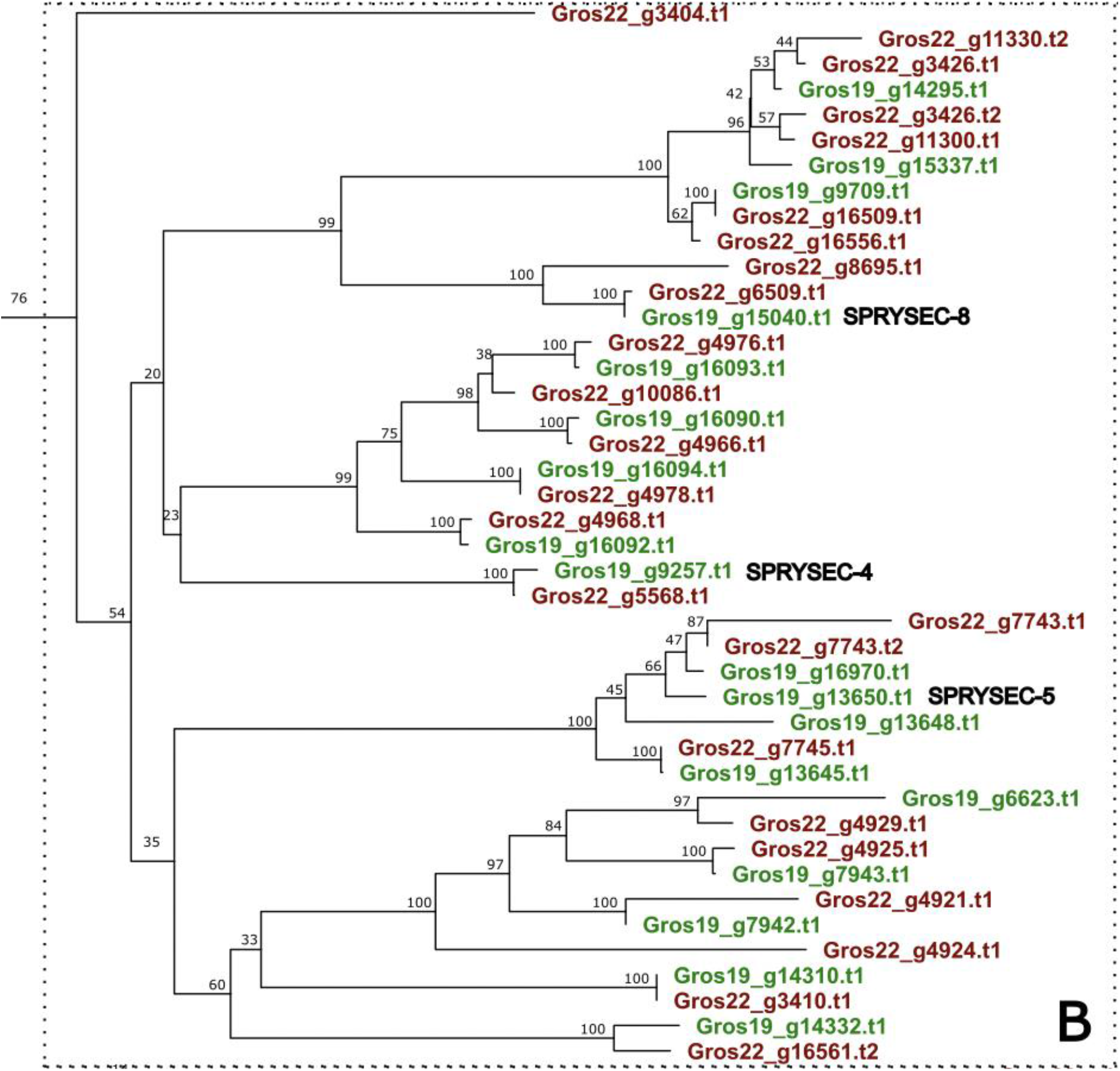
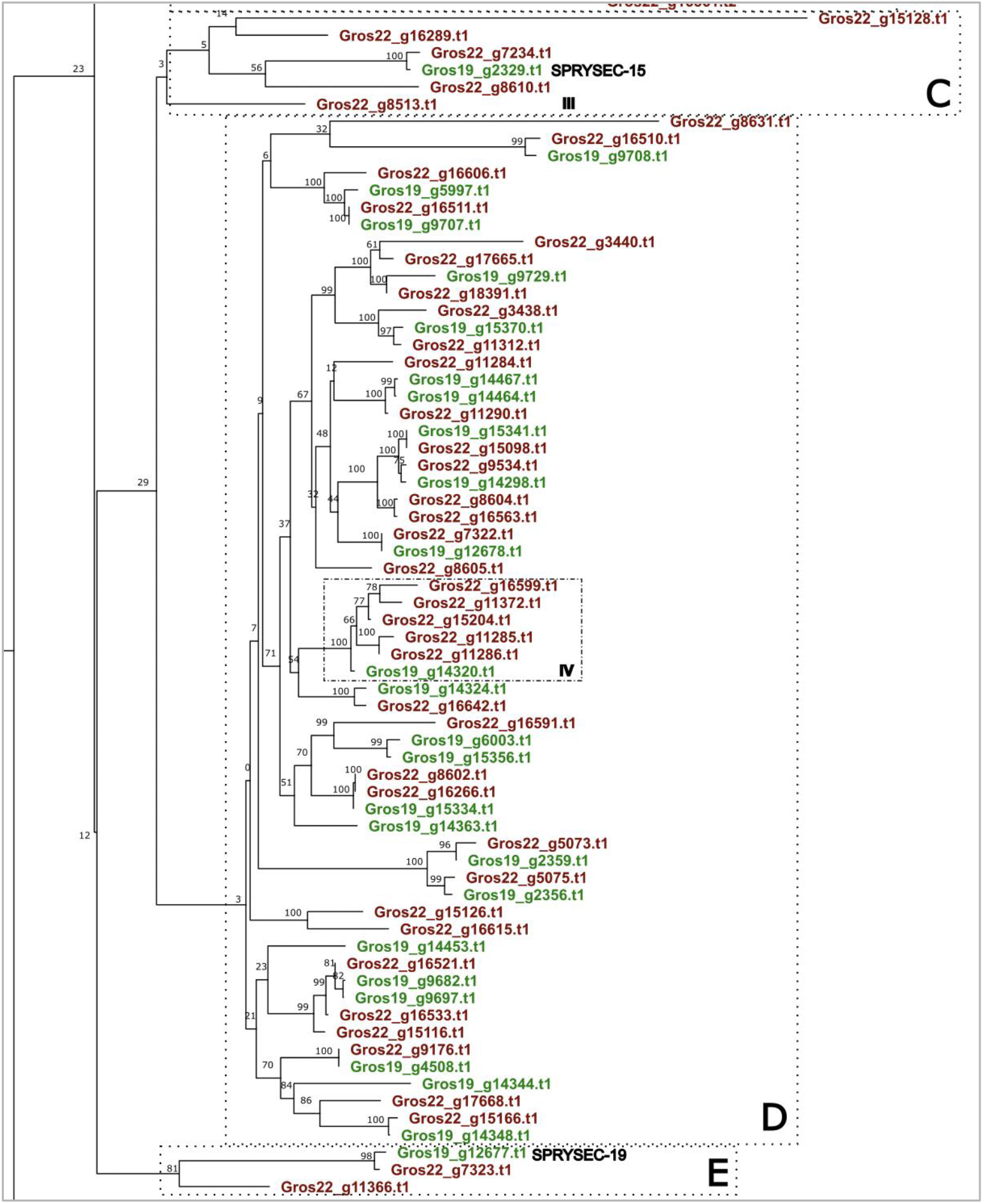
Phylogeny of SPRYSEC effector genes of both Gr-Line19 (green) and Gr-Line22 (dark red). Only SPRY proteins with a signal peptide for secretion are included A multiple sequence alignment was made using MUSCLE on the coding sequence. A phylogenetic tree was made using RAxML using a GTRGAMMA model, validated by 100 bootstrap replicates. Closest homologs to the functionally described SPRYSEC-4, SPRYSEC-5, SPRYSEC-8, SPRYSEC-18, and SPRYSEC-19 are shown.

As compared to B and D, cluster A shows the highest level of diversification. Both types of asymmetric SPRYSEC expansion were found in this cluster. Box I in Cluster A comprises a single Gr-Line22 and three Gr-Line-19 SPRYSECs. Box II exemplifies Gr-Line22 expansion, where four closely related Gr-Line22 SPRYSECs surround a single Gr-Line19 variant.

Cluster B harbours three of the SPRYSEC variants described in Ali *et al.* (2015) (SPRYSEC-4, −5 and −8), all of which seem to be represented by a single orthologous pair. As the orthologs in both lines are highly similar, we assume that these SPRYSECs are actively suppressing the host plant immune response in both lines.

Cluster C is characterized by a set of genes homologous to SPRYSEC-18 that are considerably expanded in Gr-Line22. Box III shows one Gr-Line19 copy, whereas five SPRYSEC variants represent Gr-Line22.

Cluster D unites SPRYSEC variants with a low degree of diversification. Although most Gr-Line-19 variants have a single equivalent in Gr-Line22, there are a few examples of further diversification in Gr-Line22. Box IV shows a notable example of a diversification event where a single Gr-Line19 variant has five closely related equivalents in Gr-Line22.

SPRYSEC-19, a variant that was demonstrated to suppresses programmed cell death mediated by several immune receptors (Postma *et al.*, 2012), localized in cluster E. SPRYSEC-19 was first identified in a *G. rostochiensis* Ro1 Mierenbos population (Rehman *et al.*, 2009), which is the population Gr-Line19 was originally derived from. Cluster E shows the Gr-line 22 equivalent of SPRYSEC-19 (g7323).

### D. CLE-like

The CLE-like gene family is an unusual effector family coding for prepropeptides that are delivered via the stylet of the infective J2 to the syncytial cell. For CLEs from the related cyst nematode species *Heterodera glycines* with domain structures similar to *G. rostochiensis*, it was shown that the mature propeptide comprised a nematode-specific translocation signal that facilitated the export from the developing syncytium to the apoplast (Wang *et al.*, 2020). Subsequently, the protein is cleaved outside the plant cell, and bioactive CLEs are released (Gheysen & Mitchum, 2019). Two classes of CLE-like proteins were found to be expressed in the dorsal gland of *G. rostochiensis*. Members of the *Gr-CLE-1* class showed moderate (≈ 10 fold) upregulation in early parasitic life stages (peak in parasitic J-3), whereas *Gr-CLE-4* representatives showed an over 1,000 fold in later parasitic stages (at 21 dpi) (Lu et al., 2009).

The two *G. rostochiensis* lines 19 and 22 harbour 10 and 16 CLE variants, and both lines comprise members of the *Gr-CLE-1* and the *Gr-CLE-4* class. As shown in the phylogenetic analysis (Fig. 8A), members of class *Gr-CLE-4* show little variation among each other, while *Gr-CLE-1*s show a higher level of diversification. As compared to Gr-Line19, the number of *Gr-CLE-4* variants had doubled from four to eight in Gr-Line22. On the contrary, each of the Gr-Line19 representatives of *Gr-CLE-1* had a single homolog in Gr-Line22. In Fig. 8A, clusters A and B include four Gr-Line22 variants with no immediate ortholog in Gr-Line19. In cluster C, a Gr-Line19 variant is present that deviates substantially from the closest Gr-Line22 orthologous sequence. Cluster D contains an example of a homologous gene pair, with a tentative duplication in Gr-Line22.

**Fig. 8A.**
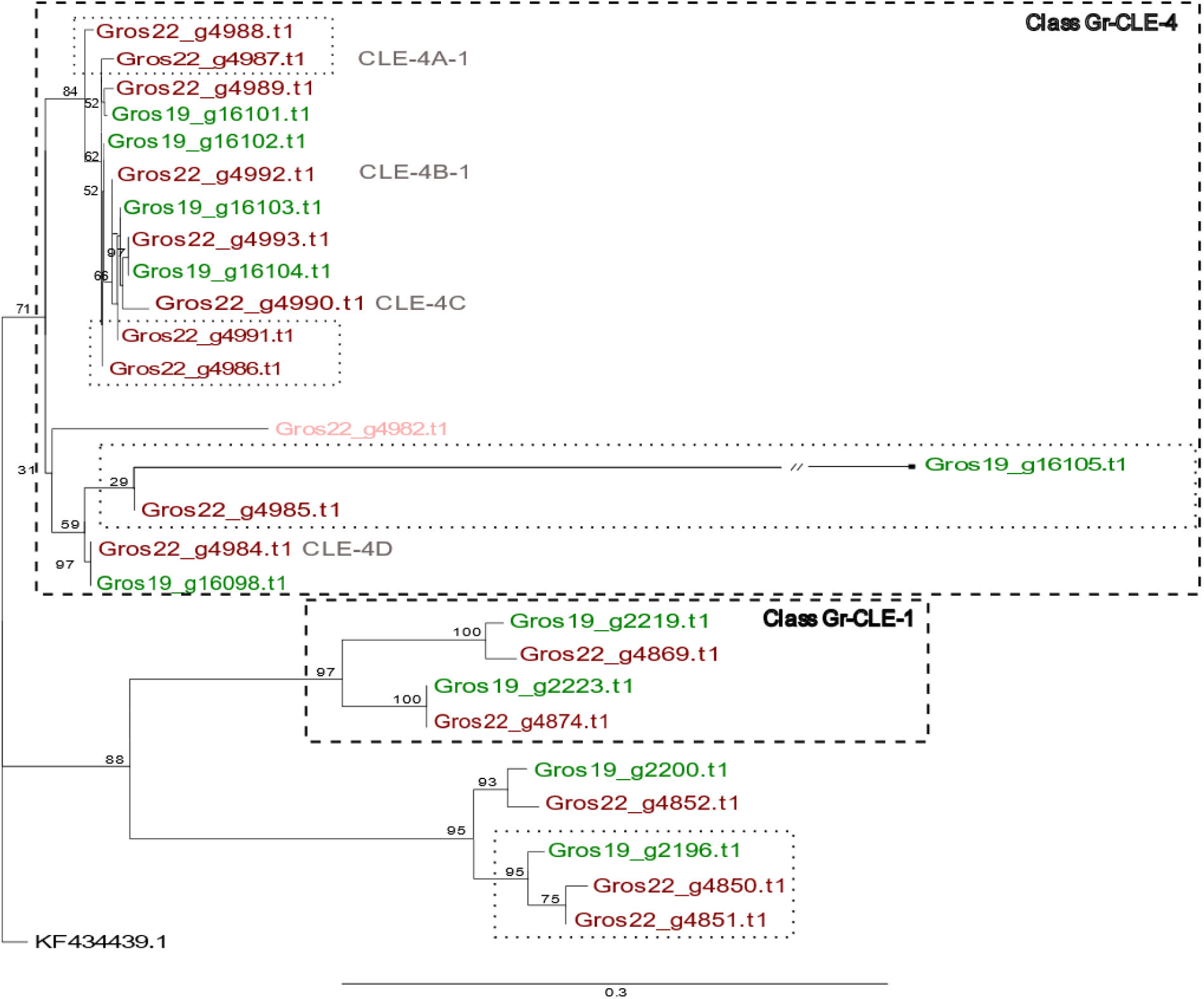
Phylogeny of CLE effector genes of both Gr-Line19 (Green) and Gr-Line22 (dark red). A multiple sequence alignment was made using MUSCLE on the coding sequence. A phylogenetic tree was made using RAxML using a GTRGAMMA model, validated by 100 bootstrap replicates. Lighter shades of green or red indicate effector variants that lack a signal peptide for secretion. Genes belonging to the functional classes GR-CLE-1 and Gr-CLE-4 are labeled with dashed boxes.

**Fig. 8B.**
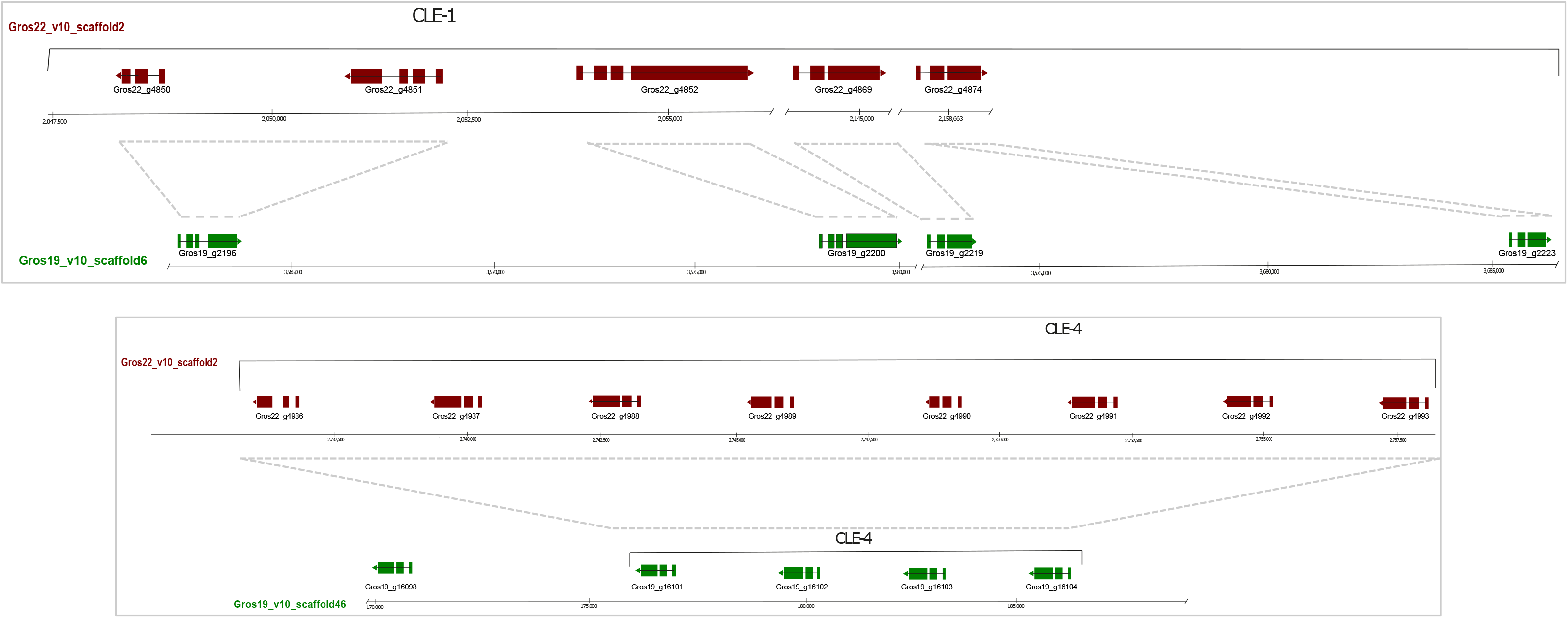
Spatial organization of CLE functional classes CLE-1 and CLE-4 for both Gr-Line19 (Green) and Gr-Line22 (Dark red)

In addition to sequence similarity within the Gr-CLE function classes, there is also a high degree of physical clustering (Fig. 8B). The Gr-CLE-4 variants are all located adjacent to each other, and not interspersed by any other gene. Based on this remarkable physical organization, we hypothesize that one or more duplication events in this region underlies the copy number difference of Gr-CLE-4 effectors (n = 4 in Gr-Line19; n=8 in Gr-Line22).

## Discussion

Due to specific biological characteristics of plant-parasitic nematodes, host plant resistances tend to be a remarkably durable means to manage this category of soil-borne pathogens. The main challenge is the actual developing and breeding resistant host-plant varieties. As the genetic basis for virulence in plant-parasitic nematodes is unknown, breeding for resistance can only be done on a trial-and-error basis. The whole process is, therefore, inefficient, and thus time consuming and expensive. The availability of molecular-based pathotyping methods of plant-parasitic nematode populations would allow for the design of more targeted resistance. Here we concentrated on two *Globodera rostochiensis* inbred lines, Gr-Line19 and Gr-Line22, with distinct pathotypes (Janssen *et al.*, 1990). Resulting from a single male-female crossing by Janssen et al. (1990), each of these lines’ genomic background is small, with a maximum of 4 haplotypes per locus. These small genomic backgrounds signifcantly simplify the generation of high-quality reference genome sequences, which has been a challenge for sexually reproducing plant-parasitic nematodes in the past. Therefore, we expect that the reference genome sequences of Gr-Line19 and Gr-Line22 are a more accurate representation of the *G. rostochiensis* genome, making the process of molecular pathotyping a step closer. Furthermore, long-read sequencing technology allowed us to generate reference genomes about 24 fold less fragmented than the current reference genome (Eves-van den Akker *et al.*, 2016). This higher continuity made it possible to pinpoint the physical distribution and the diversification in a way that was not possible with the highly fragmented JHI-Ro1 reference genome sequence. Four effector families that, together with other effectors, define this potato cyst nematode’s pathogenicity were explicitly studied in detail.

One of the main technical challenges we tried to overcome was generating high-quality reference genome sequences of a highly heterozygous nematode species. In terms of assembly sizes, we see a comparable size to the *G. rostochiensis* JHI-Ro1 genome (Eves-van den Akker et al., 2016) as well as to the estimated *G. rostochiensis* genome size (Grisi *et al.*, 1995). Therefore, it is likely that the high levels of heterozygosity did not negatively impact the assembly by the presence of haplotigs (Roach *et al.*, 2018). Another possibility is that the presence of many haplotypes negatively influenced the fragmentation of the assembly. Due to more variation in the bases, it might have been more challenging to combine more contigs into scaffolds. To further reduce the number of scaffolds, possibly to a chromosome level, it might be advantageous in the future to supplement long-read sequencing with other techniques such as optical mapping (Deschamps *et al.*, 2018; Field *et al.*, 2020).

We furthermore assessed the effect of generating a genome assembly of a highly inbred line instead of a regular population. A comparison was made between SNPs’ zygosities called on short-read data and found that Gr-Line19 had a 10% higher proportion of homozygous SNPs than the JHI-Ro1 population. This suggests that there is indeed a smaller number of haplotypes present in the inbred-lines than in the JHI-Ro1 population. Since more than 50% of the called SNPs in Gr-Line19 are still heterozygous, it seems reasonable to assume that the measured heterozygosity levels provide a more realistic picture of the heterozygosity that is present in an individual. Which is in line with previous findings in the cyst nematode *Heterodera glycines* by Ste-Croix *et al.* (2021), who described that individuals can have mixing levels of zygosity.

A more detailed analysis of four effector families for which at least a subset of members are known to be expressed in the dorsal gland of nematodes during feeding site induction or maintenance revealed dozens of novel potential virulence-associated variants.

To some extent, our starting point was comparable to the approach taken by (Bekal *et al.*, 2008). Within the soybean cyst nematode *Heterodera glycines* subsets of populations with comparable pathogenicity have been defined based on their multiplication characteristics on a set of seven soybean indicator lines. Populations that shared multiplication characteristics were coined ‘HG types’ (Niblack et al., 2002). Subsequently, Bekal et al. (2008) used two inbred lines that were either avirulent (‘TN10’; HG type 0)) or virulent (‘TN20’; HG type 1, 2, 3, 4, 5, 6, 7). 454 micro-bead sequencing of these indicator lines resulted in the generation of tens of millions of short reads (110 – 120 bp), which allowed for whole-genome comparative analysis. These efforts resulted in 239 homozygous SNPs between TN20 and TN10 (Bekal *et al.*, 2008). Although the relationship between these SNPs and pathogenicity is unclear, these SNPs could be considered one of the first molecular markers for pathogenicity in cyst nematodes. Here we took it one step further, by identifying copy number variation that might serve as potential pathotype specific molecular markers. Copy number variation is relevant, as it has been linked to virulence in various pathogens (Brynildsrud *et al.*, 2016; Zhao & Gibbons, 2018) including plant parasitic nematodes (Castagnone-Sereno et al., 2019).

For potato cyst nematodes, Folkertsma *et al.* (1996) used AFLP assays (Vos *et al.*, 1995) to characterize pathotypes of the potato cyst nematodes *G. rostochiensis* and *G. pallida.* Almost 1,000 marker loci were employed to genotype populations of both potato cyst nematode species. These analyses revealed genetic markers that can distinguish between the *G. rostochiensis* pathotypes Ro1, Ro3 and Ro4, while such loci appeared to be absent for the *G. pallida* pathotypes Pa2 and Pa3. In a more extensive approach focussing on *G. rostochiensis* only, Mimee *et al.* (2015) employed a restriction enzyme-based genotyping-by-sequencing approach. The genotypic characterization of 23 populations, covering all five pathotypes, revealed a clear distinction between pathotypes Ro1 and Ro2 on the one hand, and Ro,3, Ro4, and Ro5 on the other. Moreover, their analyses seemed to demonstrate intra-pathotype variation within Ro1. However, it is noted that with 14 populations from 9 different countries, Ro1 was overrepresented in this research.

The first reference genome for *G. rostochiensis* was published by Eves-van den Akker *et al.*, (2016), and - in conjunction with this - the intra-species variation regarding members of known effector families was mapped. All five *G. rostochiensis* pathotypes were represented in this study, and whereas homozygous molecular markers to discriminate between Ro4 and Ro5 could be identified, this was not possible for the remaining three pathotypes. Moreover, this research confirmed the large genotypic diversity of populations that are all labeled Ro1, indicating that there are many possible genotypes that yield a similar Ro1-like virulence. Here it might be mentioned that Ro1 and Ro4 share the inability to parasitize potato genotypes that harbor the *H1* resistance gene from *Solanum tuberosum* ssp. *andigena* CPC 1673. Moreover, the H1 resistance genes have been introgressed in most commercial potato varieties, and potato cyst nematode populations worldwide have been exposed to these resistance genes likely including the ones characterized by Eves-van den Akker *et al.* (2016). So, although these pathotypes share their avirulence concerning the *H1* gene, they belong to another *G. rostochiensis* genotype and differ significantly in intra-pathotype variation.

Hence, as a starting point, we used pathotypically characterized inbred lines from which we generated new reference genome sequences. On this basis, complete effector families could be mapped and compared. In essence, the make-up of effector families in lines with distinct pathogenic characteristics could vary because of (1) non-synonymous variants in sequence in a given set of effector genes and/or (2) effector gene loss or gain (3) quantitative variation in expression levels due to SNPs in the promotor region (4) quantitative variation in expression levels due to copy number variation. The balance between these two (dependent) sources of variation varies in a pathogen-dependent manner. The genome-wide comparison of three *Microbotryum* species parasitizing distinct Caryophyllaceae allowed Beckerson *et al.* (2019) to define the secretomes of the individual species. Their analyses revealed that host specificity was explained by rapid changes in effector genes rather than by variation in the effector copy numbers. With a similar underlying question, Qutob *et al.* (2009) investigated two effector genes families of *Phytophthora sojae,* Avr1a and Avr3a in a range of races. The presence of multiple copies of nearly identical genes on the Avr1a and the Avr3a locus was suggested to contribute to the fitness of these races, and races with distinct pathogenicities were characterized by variations in effector gene numbers. These examples demonstrate that both sources of variation can generate differences in pathogenicity among plant pathogens. Here we specifically focussed on effector gene loss and gain effects, and observed that both events happen in the avirulent Gr-Line19 as well as the virulent Gr-Line22. Previous studies show that, at least in potato cyst nematodes, single nucleotide polymorphisms are also related to virulence (e.g. Sacco *et al.* (2009)), which indicates that both types of genomic variance are relatable with virulence.

In case of the tropical root-knot nematode *Meloidogyne incognita,* Castagnone-Sereno et al., (2019) tried to pinpoint the genetic basis of avirulence and virulence with regard to the tomato resistance gene *Mi-1.2.* Genome-wide characterization of two pairs of avirulent and virulent lines revealed 20 gene families that all showed a lower number of copies in both virulent *M. incognita* lines. It is noted that the 20 families included pioneers and household genes, and not known effector families. Hence, although a lower copy number per gene family was associated with virulence, this research did not identify gene loss events that could be causally related to virulence.

We separately considered the dorsal esophageal gland-expressed effectors that are thought to be involved in immune response suppression and feeding site induction, and the subventral gland-expressed effectors that are active during plant penetration. Concerning dorsal gland-expressed effector families, *G. rostochiensis* Gr-Line19 harboured on average 14 genes per effector family, while on average, 19 members were identified per effector family in Gr-Line22, a homozygous virulent line with regard to the H1 resistance gene. Two exceptions to this overall increase in effector copy number should be mentioned; GSS appeared to be absent in Gr-Line22, while lower copy numbers were observed for Hg-GLAND6. GSS is a peculiar example as this was the first example of a cyst nematode effector family that arose from neofunctionalization of a household gene, glutathione synthetase (Lilley *et al.*, 2018). Two expansion waves of GS-like genes are described, and only in the second wave GS-like genes with a signal peptide for secretion arose. This clade 3 comprises cyst nematode effectors. It should be noted that *GSS* genes in this clade share only 36% protein identity. The thresholds used in this study to identify *GSS* family members were presumably too strict to identify more GS-like members (and even none in Gr-Line22).

The effector families expressed in the subventral glands that are included in this study showed less expansion than the dorsal gland specific families, with only small differences in copy numbers between the two lines. Strikingly, a substantial number of genes belonging to this category lack a signal peptide presence. Since many of these genes (e.g., glycoside hydrolases, pectate lyases) code for cell-wall degrading or modifying enzymes, the proteins would have to be secreted to make physical contact with the plant in order to perform their function. One hypothesis could be that these genes are, in fact, pseudogenes. However, this seems unlikely as manual inspection showed that most of these SP lacking genes show a RNAseq signal (results not shown). Whereas ample RNAseq data allowed for an accurate prediction of the intron-exon structure, the transcription start site is more difficult to predict without additional experimental data. If transcription start sites were misplaced, we could have missed a preceding signal peptide. Alternatively, it could be that this cyst nematode genuinely harbours effector variants without apparent signal peptide similar to the invertases identified in *Meloidogyne incognita.* These effectors were suggested to be acquired at a late stage during cyst nematode evolution (Abad *et al.*, 2008).

Phylogenetic analysis of effector families as presented here takes along both effector diversification and effector loss and gain. These data clearly demonstrate that the balance between both sources of variation differs per effector family. Whereas effector family 1106 showed overall little copy number variation, SPRYSECs showed a 65% increase in copy number in Gr-Line22, and in case of the GLAND5 family significant diversification was accompanied by a sharp increase of the copy number in Gr-Line22. Other population genetic studies on plant-pathogenic fungi and oomycetes showed exclusively low (Talas & McDonald, 2015) or high (Flier *et al.*, 2003) levels of diversification between effector genes. We are not aware of other plant pathogens for which such drastic contrasts in diversification pattern between effector families were described.

Because of its extreme level of physical clustering of CLE effectors in both *G. rostochiensis* inbred lines we investigated its genomic organisation in more detail. Potato cyst nematodes produce and secrete mimics of plant CLEs. Plant CLEs are signalling components that were shown to be conserved in both Arabidopsis and potato roots (Lu *et al.*, 2009). Among *Globodera* CLE genes two functional classes are distinguished, CLE1 and CLE4. The main difference between these classes is the composition of CLE peptides that are present as small cleavable units separated by small spacers at the protein’s C terminus. Mitchum *et al.* (2012) described a single CLE1 representative, and here a second potential CLE1 variant is identified in both *G. rostochiensis* lines (Fig. 8a). This second variant has a domain structure similar to GrCLE1 (Supplemental Fig. 1), and the conservation of the domain structure makes it plausible this variant has a CLE1-like function. Notable is the putative duplication event of Gr-CLE4 genes in Gr-Line22. Gr-CLE-4 genes are highly conserved, even between pathotypes and we assume that this duplication event might result in a higher production of GrCLE4 peptides. A dose effect for a nematode effector was previously reported for the 32E03 effector of the beet cyst nematode *Heterodera schachtii* (Vijayapalani et al., 2018). So our finding might suggest that the virulent *G. rostochiensis* line 22 might exert a stronger CLE4 peptide-based effects on its host.

### Outlook / future prospects

Molecular pathotyping is an essential element in durable disease management. After all, this will allow breeders to use host plant resistances in a targeted way, and it allows farmers to make a more informed decision which potato variety to grow in the field. The existing pathotyping system for *G. rostochiensis* classifies populations into five pathotypes (Ro1-Ro5) on the basis of their relative multiplication rates on a number of *Solanum* differentials, and this systematic was used as starting point for the generation of new pathotyping platform. By generating high quality reference genomes from two pathotypically-distinct inbred lines, we were able to generate broad overviews of effector families including their diversification and spatial organisation. On the basis of a selection of four effector families, dozens of effector variants could be pinpointed that were unique for either of the two inbred lines Gr-Line19 (Ro1) and Gr-line22 (Ro5). Once these data are supplemented by re-sequencing data from well-characterized *G. rostochiensis* field populations, comparative effectoromics would be within reach. Comparative effectoromics will provide a fundament for our understanding of compatible and incompatible host-nematode interactions as well as for a new, biologically insightful pathotyping scheme as a basis for the durable use of host plant resistances.

## Acknowledgements

We thank John T. Jones for his expertise in the assembly of plant parasitic nematode genome sequences.

## Contributions

SvdE performed the DNA extraction and library preparations. PT and JvS designed the assembly pipeline. JvS generated the genome assemblies. JvS performed the comparative genomics/effectoromics and phylogenetic analyses. JvS and JH wrote the manuscript; all other authors co-commented on the manuscript.

## Ethics declarations

Ethics approval was not needed for this study.

